# Changes and composition of microbial community during aerobic composting of household food waste

**DOI:** 10.1101/2021.03.12.435218

**Authors:** Zhihua Jin, Tong Lu, Wenjun Feng, Qinchao Jin, Zhige Wu, Yu Yang

## Abstract

In order to explore the effect of added bacteria on microbial community and determine the dominant bacteria in the aerobic composting process of household food waste (HFW), two groups of HFW composting experiments were conducted for 30 days. The final degradation rates of the two groups were 69.95% (group A, natural composting) and 73.52% (Group C, composting with added bacteria), respectively. 16S rRNA high-throughput sequencing was used to analyze the changes of microbial community in the composting process. As the result displays, at the classification of phylum level, the relatively abundant bacteria of two groups were *Firmicutes, Proteobacteria* and *Cyanobacteria*. At the classification of genus level, *Pediococcus* is the dominant bacteria of group A, which performed better in maintaining the microbial community stable in the later stage of composting, while *Weissella* accounted for a relatively large proportion of group C and behaved well in promoting the final degradation rate of composting. The proportion of *Ochrobactrum* in the early stage of group C is considerable and it is closely related to the removal of odour during composting. The relative abundance of added bacteria was always in a rather low level, suggested that the way they affect the composting process is to change the proportion of dominant bacteria in different stage of composting. This experiment provides an important reference for improving the microbial degradation efficiency of HFW.

**IMPORTANCE:** In recent years, food waste has gradually become a global problem, the annual waste of food is as high as 1.3 billion tons. FW, especially HFW, as a high content of organic matter waste, has a pretty good recycling value. So how to deal with and recycle it efficiently, quickly and conveniently becomes more and more important. Among many treatment and recovery methods, microbial treatment (including anaerobic digestion and aerobic composting) has gradually become a research hotspot due to its advantages of low pollution and low energy consumption, and microorganisms play a crucial role in these process.

In this study we use 16S rRNA high-throughput sequencing method to explore aerobic composting of HFW. The purpose of this study is to find out the dominant bacteria which can improve the degradation efficiency, remove the odor and prolong the treatment cycle, and then provide further theoretical reference for future HFW related research.

## INTRODUCTION

With the improvement of people’s living standards, the proportion of food waste (FW) is also increasing year by year, being considered to be one of the largest groups of organic solid waste in the world, and its generation rate is rising steadily (1,2). FW mainly comes from household kitchens, restaurants, canteens, and food processing industries (3). Among them, household food waste (HFW) is getting much attention owing to its high organic content and excellent source of value added products (4). About 1.3 billion tons of edible food are lost or wasted every year in the world (5), So the effective and harmless treatment of FW has always been the focus of global attention (6,7).

At present, the treatment methods of FW mainly include direct crushing, dehydration, chemical treatment, incineration and microbial treatment etc(8). Most of these methods consume high energy and even cause secondary pollution to the environment except microbial treatment (9). Therefore, microbial treatment of FW has become a research hotspot at home and abroad. Microbial treatment methods include anaerobic digestion and aerobic composting, both are the process of decomposing complex organic matter into small molecule organic or inorganic matter by the metabolism of microorganisms (10). This process can be realized not only by indigenous microorganisms naturally growing in FW, but also by bacteria added during composting (11). Anaerobic digestion and aerobic composting have good resource recovery properties, and have the ability to produce high value-added products (12). Thus, microbial treatment methods present a good impetus and vast potential for future development (13). For instance, Quashie Frank Koblah, et al. use a continuous stirred microbial electrolysis cell (CSMEC) with anaerobic digestion to deal with FW and produce methanation and bioelectricity. The final COD removal of CSMEC surpassed 92% with OLR (organic load rate) ranging from 0.4 to 21.31 kg COD/m^3^·d,and *Firmicutes, Proteobacteria*, and *Euryarchaeota* were the dominant phyla observed (14). The natural composting method was implemented by Tran Huu Tuan et al. to study the degradation effect of high concentration of dietyl terephthalate (DOTP) FW by natural microorganisms proliferating during composting. The total degradation efficiency of DOTP reached 98%,and *Firmicutes* was the most dominant at the phylum level, followed by *Proteobacteria* and *Bacteroidetes* (15).Huang WY et al. determined *Proteobacteria, Actinobacteria, Bacteroidetes* and *Firmicutes* as the main phyla during aerobic FW co-composting degradation of highly PCDD/F-contaminated field soil (16). Wang et al. reported that the dominant phyla of the community structure in fed-batch composting were *Firmicutes*, *Proteobacteria*, *Bacteroidetes*, and *Actinobacteria* by high-throughput sequencing (17). Composting is an effective process for the valorisation of HFW into a stable and nutrient-enrich biofertiliser (18), in which a variety of microorganisms play a critical role (19). HFW is a kind of non-uniform material with high content of water, oil, salt and cellulose, etc (20). Gaseous emissions, such as methane (CH_4_), nitrous oxide (N_2_O), and ammonia (NH_3_) which hinder composting application for food waste treatment (21), are inherent by-products of the composting process regardless of the initial organic material or process condition (22). These particular characteristics will affect some aspects of the composting quality and process. As microorganisms play a key role in the composting process, inoculating microorganisms with specific functions may have a positive effect on the composting process and improve the composting quality. So far, there are few studies on the changes of microbial community in the process of HFW composting. In this study, 16SrRNA high-throughput sequencing was used to analyze the microbial community changes and composition in different stages of natural composting and composting with added bacteria, aiming to find out the dominant bacteria which can remove odor, improve the degradation rate and prolong the fermentation cycle in the composting process, so as to provide a theoretical reference for the future aerobic composting process of HFW.

## RESULT AND DISCUSSION

### Analysis of degradation rate and viable count

Table 1 shows the changes of related parameters in different stages of composting process. The final degradation rates of group A and group C were 69.95% and 73.52% respectively. From the perspective of degradation rate, there was no significant difference between group A and group C. However, from the perspective of specific composting process, during stage 1-3, the odor produced by group C was significantly less than that by group A, and the composting residue of group C became looser and drier, which is helpful for the follow-up treatment of HFW. In terms of the proportion of composting residue in the machine volume, group C showed a good degradation efficiency in each stage of composting process, being consistent with the result of final degradation rate. From the change of pH in the composting, with the continuous addition of HFW, the pH values in the machine of two groups decreased gradually, while the pH value increased during 10 days naturally storage after the composting process, indicating that the composting process of HFW was an acidic process, which may be more suitable for the growth of some acidophilic strains. The total number of viable bacteria in group A was larger than that in group C. During the process from stage 1 to stage 2, the number of viable bacteria in group A decreased while that in group C increased, which may be as a result of the inhibitory effect between the added bacteria and the bacteria naturally growing during the composting process. In the later stage, the microbial community gradually stabilized and the number of viable bacteria began to increase, and then gradually decreased. The number of viable bacteria in the two groups decreased slowly from stage 3 to stage 5. The reason for this phenomenon may be that the decline of pH value changed the composting environment of the microbial community, at the same time, the increase of composting residue in the machine volume leads to the decrease of oxygen content in the machine, leading to the decrease of aerobic composting bacteria. These two factors may be the key points of the decreasing of viable bacteria during stage 3-5.

**Table 1:** Changes of related parameters in different stages of composting process

| Samples | Composting<br>time (d) | Viable count<br>(CFU/g) | pH | Percentage of composting<br>residue in machine volume |
| --- | --- | --- | --- | --- |
| A1 | 1 | $6.97 \times 10^{11}$ | 6.21 | 7.94% |
| A2 | 7 | $2.63 \times 10^8$ | 5.95 | 18.24% |
| A3 | 15 | $2.23 \times 10^6$ | 5.56 | 32.35% |
| A4 | 20 | $1.06 \times 10^6$ | 5.32 | 41.17% |
| A5 | 30 | $8.66 \times 10^5$ | 5.87 | 40.58% |
| C1 | 1 | $2.21 \times 10^6$ | 6.35 | 7.06% |
| C2 | 7 | $7.63 \times 10^7$ | 6.13 | 16.18% |
| C3 | 15 | $4.33 \times 10^5$ | 5.88 | 30.30% |
| C4 | 20 | $2.33 \times 10^5$ | 5.57 | 39.12% |
| C5 | 30 | $7.33 \times 10^4$ | 5.96 | 38.53% |

### Analysis of sample sequencing data

As shown in Table 2, after obtaining the original sequence data of different samples by Illumina high-throughput sequencing with optimization and statistics, the final number of effective sequences of 10 samples was 1286075, and the range of effective sequences of each sample was 121536-133111, of which 647227 were obtained from 5 samples in group A and 638848 were obtained from 5 samples in group C. The average sequence length was 422.95 bp. Based on 97% sequence homology, a total of 6626 OTUs were obtained, including 4144 OTUs in group A, and 3282 OTUs in group C. After filtering out the OTUs containing Chloroplast, 5949 OTUs were obtained, including 3804 OTUs in group A and 2945 OTUs in group C. The results of Venn diagram (Fig. 1) showed that there were 800 OTUs shared by these two groups, accounting for 19.31% of group A and 24.38% of group C, respectively. By comparison, it was found that there was a significant difference in OTUs between the two groups, which might be due to the effect of the added bacteria on the composition and structure of microbial community.

**Table 2:** Statistics of optimized effective sequences and OTUs in each sample

| Sample | optimized effective sequences | OTUs | OTUs (without<br>Chloroplast) |
| --- | --- | --- | --- |
| A1 | 131796 | 388 | 387 |
| A2 | 128788 | 460 | 459 |
| A3 | 130073 | 1076 | 835 |
| A4 | 132226 | 1118 | 1100 |
| A5 | 124344 | 1349 | 1270 |
| C1 | 133111 | 1300 | 1300 |
| C2 | 132463 | 565 | 565 |
| C3 | 124660 | 1624 | 1620 |
| C4 | 127078 | 824 | 717 |
| C5 | 121536 | 622 | 396 |

**Figure 1.**
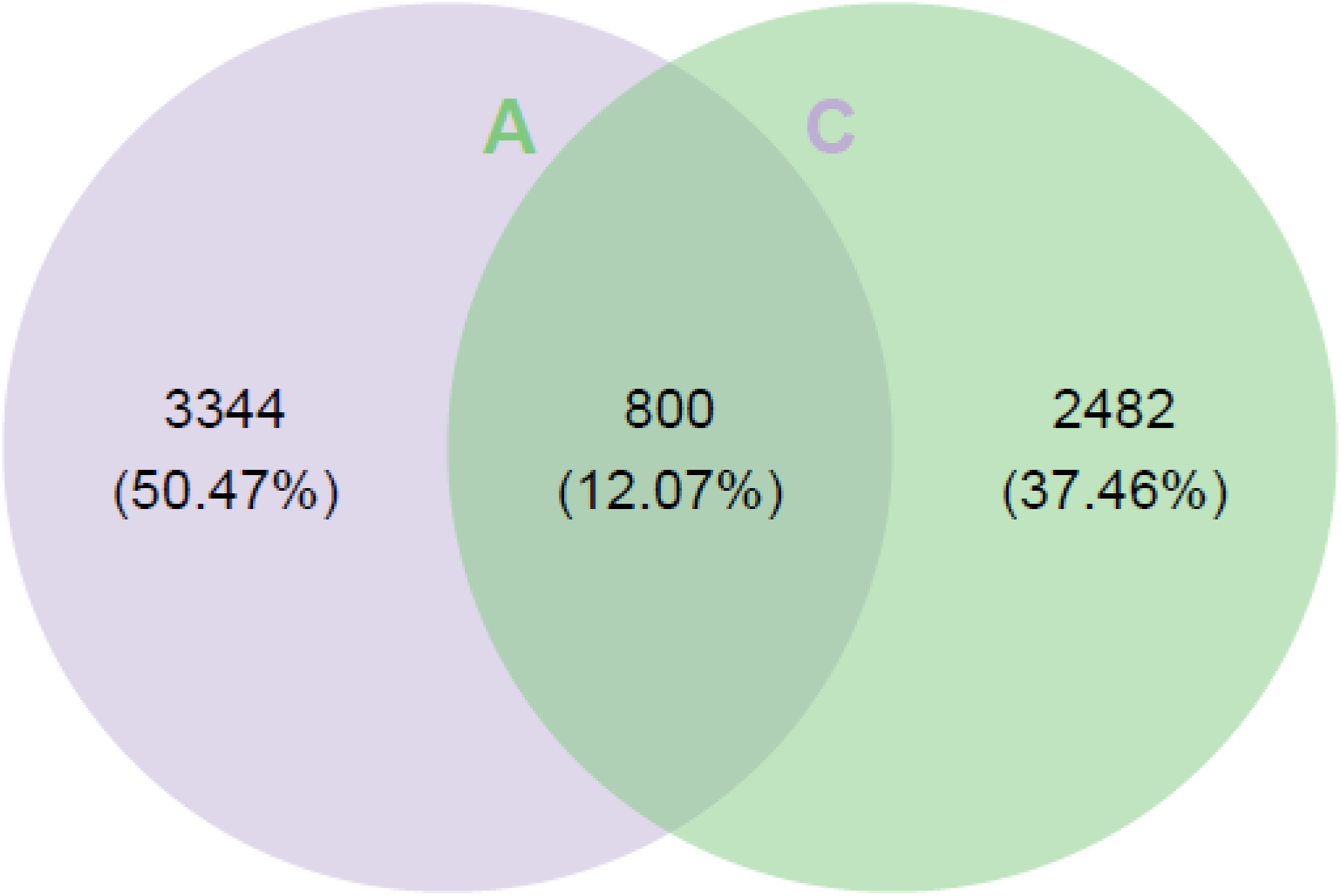
Venn graph analysis based on OTU number

### Analysis of alpha diversity index of samples

Single sample diversity analysis (alpha diversity) reflects the abundance and diversity of microbial community. The alpha diversity index of each group is shown in Table 3. The results showed that the Chao1 index and Shannon index of group A increased steadily, and reached the maximum value in A5. The Simpson index of group A reached 0.7867, 0.8860 and 0.8823 in A3, A4 and A5, respectively, indicating that the dominant bacteria of group A could be in a stable position for a long time in the middle and late composting period. The Simpson index could still maintain a high level in A5, indicating that the microbial community of group A has a good ability to maintain self-stabilization, which is very important for the extension of composting cycle. In group C, the Chao1 index and Shannon index of the microbial community fluctuated greatly, decreased firstly and then increased, and finally reached the maximum value of 1632 and 6.61 respectively in C3. The Chao1 index and Simpson index decreased significantly in C4 stage, which indicated that the microbial community abundance decreased greatly and the dominant strains maintained stable only for a short time. The Chao1 index and Simpson index in C5 stage continued to decline, indicating that the ability of group C to maintain community stable was not as good as that of group A. The sequencing coverage of samples in each group was above 0.99, indicating that the sequencing had high coverage of bacteria in samples and suitable sequencing depth, which could meet the needs of bacterial diversity analysis in each group. Fig. 2 A and B show the changes of coverage index and Shannon index with the increase of sequencing depth, respectively. It can be seen from the Fig. 2 that the coverage index and Shannon index in each group of samples gradually increased and tend to be flat with the increasing of sequencing depth, which further indicates that the sequencing results can more comprehensively reflect the diversity information contained in these samples.

**Table 3:** Alpha diversity index in each group

| Sample | Chao1 index | Shannon index | Simpson index | Coverage index |
| --- | --- | --- | --- | --- |
| A1 | 390 | 2.91 | 0.8172 | 0.999824 |
| A2 | 481 | 2.22 | 0.6058 | 0.999341 |
| A3 | 1119 | 3.68 | 0.7867 | 0.998544 |
| A4 | 1146 | 5.01 | 0.8860 | 0.998834 |
| A5 | 1380 | 5.31 | 0.8823 | 0.998801 |
| C1 | 1346 | 2.64 | 0.3926 | 0.998469 |
| C2 | 607 | 1.86 | 0.4941 | 0.999011 |
| C3 | 1632 | 6.61 | 0.9585 | 0.999083 |
| C4 | 848 | 4.11 | 0.7467 | 0.998987 |
| C5 | 650 | 3.36 | 0.7235 | 0.999132 |

**Figure 2(A).**
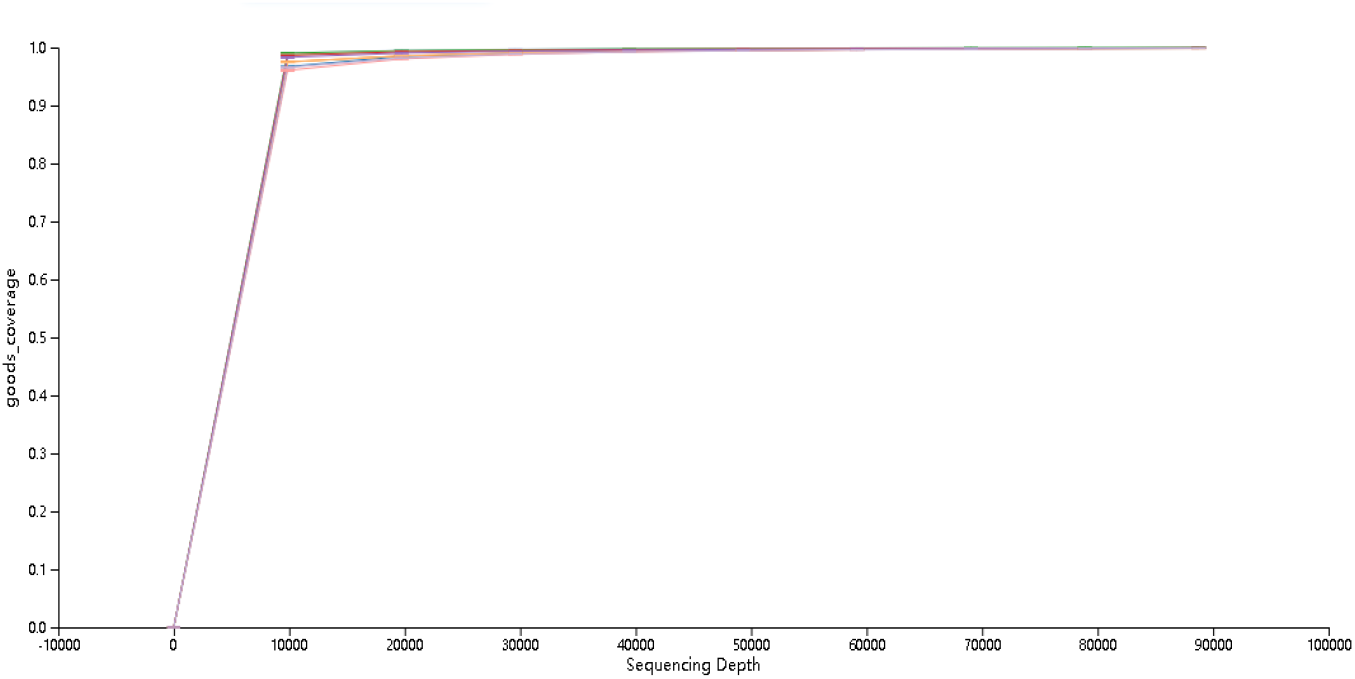
Coverage-Sequencing Depth Curve

**Figure 2(B).**
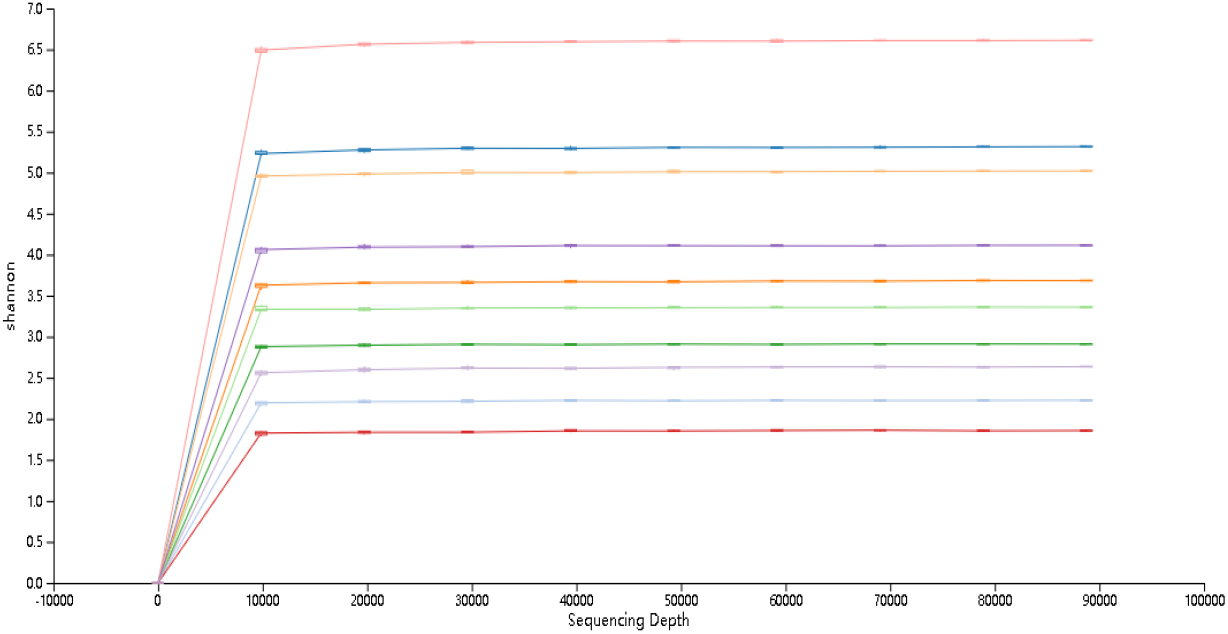
Shannon-Sequencing Depth Curve

### Analysis of bacteria composition during composting process at phylum level

Fig. 3 shows the bacteria composition in different stages at the level of phyla classification. The main bacteria phyla in stage1 is: A1: *Proteobacteria* (98.81%), *Firmicutes* (0.64%); C1: *Proteobacteria* (90.03%), *Firmicutes* (5.86%), *Actinobacteria* (2.66%). The relative abundance of *Proteobacteria* in the two groups of samples in stage1both exceeded 90%, indicating that *Proteobacteria* is in an absolutely dominant position in both A1 and C1. Many bacteria of *Proteobacteria* are pathogenic (23),such as *Cronobacter* (24), *Burkholderia* (25,26) and *Ochrobactrum* (27). The main bacteria phyla in stage2 is: A2: *Firmicutes* (96.23%), *Proteobacteria* (3.07%); C2: *Firmicutes* (96.60%), *Proteobacteria* (2.62%). In stage 2, the type and proportion of bacteria phyla in A2 and C2 periods were almost the same and the proportion of *Firmicutes* rose rapidly from stage 1 to stage 2 and replaced *Proteobacteria* as the dominant bacteria phyla. The phylum *Firmicutes* includes Gram-positive bacteria with rigid or semi-rigid cell walls that are predominantly from the genera *Bacillus, Clostridium, Enterococcus, Lactobacillus* and *Ruminicoccus* (28). Moreover, many *Firmicutes* produce spores that are resistant to dehydration and extreme environments (29), which may be one of the reasons why group A had a strong ability to maintain stable in the late stage, of which the proportion of *Firmicutes* was more than 80%. The main bacteria phyla in stage3 is: A3: *Firmicutes* (66.67%), *Cyanobacteria* (26.73%), *Proteobacteria* (4.89%); C3: *Firmicutes* (49.09%), *Proteobacteria* (40.84%), *Actinobacteria* (2.70%), *Bacteroidetes* (2.70%), *Acidobacteria* (1.19%). The emergence of *Cyanobacteria* in A3 led to a decrease in the proportion of *Firmicutes*. Those bloom-forming *Cyanobacteria* may have a certain inhibitory effect on pathogenic bacteria of *Proteobacteria*, because some genera of it possess the ability to produce various toxic or bioactive secondary metabolites, e.g. toxic microcystins (MCs) and bioactive anabaenopeptins (APs) which is linked to antibacterial, fungal or cytostatic effects (30). In contrast, the proportion of *Proteobacteria* in C3 period increased greatly without *Cyanobacteria* existing, further proving that the emergence of *cyanobacteria* in A3 could inhibit the proliferation of the pathogenic bacteria of *Proteobacteria*. The main bacteria phyla in stage4 is: A4: *Firmicutes* (93.66%), *Proteobacteria* (2.34%), *Cyanobacteria* (1.90%); C4: *Firmicutes* (78.89%), *Cyanobacteria* (15.64%), *Proteobacteria* (4.64%). *Firmicutes* still occupied the dominant position in A4, while the proportion of *Proteobacteria* in C4 decreased significantly with a small amount of *Cyanobacteria* appeared, which confirmed once again the inhibitory effect of *Cyanobacteria* on pathogenic bacteria of *Proteobacteria*. The main bacteria phyla in stage 5 is: A5: *Firmicutes* (82.30%), *Cyanobacteria* (6.87%), *Proteobacteria* (5.08%), *Bacteroidetes* (2.93%), *Actinobacteria* (2.15%); C5: *Firmicutes* (54.40%), *Proteobacteria* (2.44%), *Cyanobacteria* (42.61%). The proportion of *Firmicutes* in A5 was about 30% more than that in C5, indicating that the relative abundance of dominant phyla in group A could still maintain at a high level after ten days of storage, indicating that group A had a good ability to maintain community stable.

**Figure 3.**
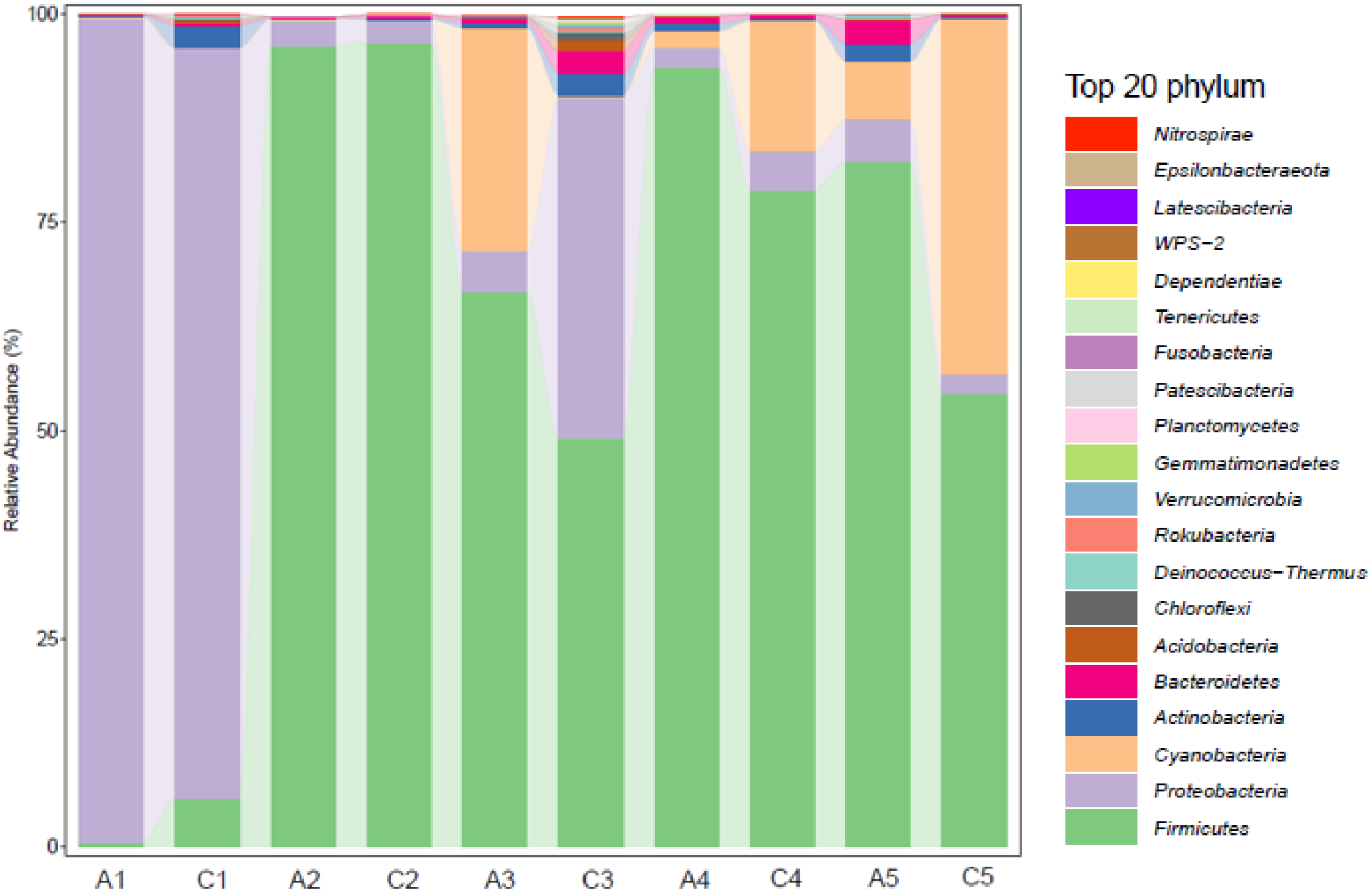
Bacteria composition at the level of phylum classification at different stages

### Analysis of bacteria composition during composting process at genus level

As shown in Fig. 4, at the level of genus classification, the relatively abundant bacterial genera in the A1 are *Cronobacter* (57.44%), unclassified_*Enterobacteriaceae* (24.75%), *Burkholderia-Caballeronia-ParaBurkholderia* (15.24%). These three genera belonging to *Proteobacteria* are all opportunistic pathogens. Now, 10 individual species of *Cronobacter* have been identified (31,32),which are associated with necrotizing enterocolitis, septicemia, and meningitis (33). *Enterobacteriaceae* family can contaminate fresh products at any stage of production either at pre-harvest or post-harvest stages (34). Many species of *Burkholderia* are opportunistic pathogens, such as *Burkholderia pseudomallei* that causes melioidosis (35) and *Burkholderia cepacia* that causes cystic fibrosis in human lungs (36,37). The relatively abundant bacterial genera in C1 is *Ochrobactrum* (84.61%), and the proportion of added bacteria is: *Bacillus* (1.89%), *Pseudomonas* (0.24%),*Paenibacillus* (0.01%).Similarly, *Ochrobactrum* also belongs to the *Proteobacteria*, and several *Ochrobactrum* spp. have been investigated for their potential to degrade xenobiotic pollutants and for heavy metal detoxification under a variety of environmental conditions (38). Moreover, the odour of group C in the early stage of composting was significantly smaller than that of group A. Therefore, *Ochrobactrum* may play a better role in degradation and deodorization of HFW in the early stage of composting. However, in the early stage, the proportion of added bacteria is less, which may be caused by the small initial addition or the inhibition of the growth of pathogenic bacteria. It can be clearly seen that the types and proportions of the dominant bacteria in the two groups in the early stage of composting are quite different, but at the level of phylum classification they all belong to *Proteobacteria*. The reason for this result may be that the presence of added bacteria affects the proportion of the dominant bacteria during the process of composting. The relatively abundant bacteria genera in the A2 are *Pediococcus* (93.12%), *Klebsiella* (1.40%), while that in the C2 are *Pediococcus* (94.82%), *Ochrobactrum* (1.29%), and the proportion of added bacteria is: *Bacillus* (0.38%), *Pseudomonas* (0.021%), *Paenibacillus*(0.02%). Obviously different from the first stage, the dominant bacteria in both A2 and C2 are *Pediococcus* belonging to *Firmicutes* and the relative abundance of them are both more than 90%. Literature review shows that *Pediococcus* can secrete pediocins which are short peptides with strong antibacterial activity, high temperature resistance, and acid resistance in the process of proliferation (39). It is noteworthy that the proportion of pathogenic bacteria in A2 and C2 drops dramatically. It is inferred that the pediocins secreted by *Pediococcus* inhibit and control the opportunistic pathogens at a low level. Meanwhile,the proportion of added bacteria is still maintained at a relatively low level, which may be due to the inhibitory effect of the proliferation of *Pediococcus*. The relatively abundant bacterial genera in A3 are *Pediococcus* (57.01%), *Bacillus* (2.53%), *Enterococcus* (1.33%), and some Chloroplast (26.73%). The proportion of *Pediococcus* in A3 decreased to a certain extent, and a small amount of Chloroplast appeared, which may be as a result of incomplete degradation of vegetables in HFW. The main genera of C3 are *Pediococcus* (20.73%), *Lactobacillus* (10.10%), *Proteus* (8.64%), *Acinetobacter* (7.65%), *Klebsiella* (7.08%) and *Oceanobacillus* (6.74%). Among these genera, *Proteus*, *Acinetobacter* and *Klebsiella* are all opportunistic pathogen. *Proteus* consists of five species and three unnamed subspecies. Among these species, *P. vulgaris* and *P. mirabilis* are most frequently linked with food contamination and food poisoning (40,41).The non-fermentative bacteria of *Acinetobacter* are opportunistic pathogens,for *instance,Acinetobacter Baumannii* has emerged as one of the most problematic common opportunistic nosocomial pathogens worldwide (42). The most common species of *Klebsiella* is *Klebsiella Pneumoniae*, a conditional pathogen that can settle in human skin and mucous membranes to cause urinary tract infections, sepsis and pneumonia in individuals with weakened immunity (43,44).The *Lactobacillus*, of which the relative abundance increased in C3, is a kind of probiotics widely existing in human intestinal tract (45), and may play an important role in inhibiting those opportunistic pathogens. The proportion of added bacteria in C3 is: *Bacillus* (2.21%), *Pseudomonas* (0.68%), *Paenibacillus*(0.11%), increased slightly, but still at a low level, which may be due to the low pH and high salt concentration. In C3, the proportion of *Pediococcus* decreased significantly, and a large number of miscellaneous bacteria appeared. The reason for this phenomenon may be that when the proportion of *Pediococcus* decreased and *Weissella* did not occupy the dominant position in the middle stage of composting, these pathogenic bacteria and miscellaneous bacteria took this opportunity to proliferate and occupy a certain proportion. As the proportion of *Weissella* increased in later stage, the relative abundance of these miscellaneous bacteria decreased sharply to a very low level. Members of the genus *Oceanobacillus* were aerobic, rod-shaped, endospore-forming, halophilic bacteria (46). The increase of *Oceanobacillus* in C3 may be due to the rising of salt concentration in the middle stage of composting. The relative abundance of bacteria genera of Stage 4 was as follows, A4: *Pediococcus* (41.39%), *Weissella* (34.97%), *Leuconostoc* (5.68%), *Bacillus* (2.08%)*, Lactobacillus* (1.55%), *Macrococcus* (1.55%); C4: *Weissella* (70.89%), *Leuconostoc* (3.07%), *Acinetobacter* (1.16%), and added bacteria of *Bacillus* (0.46%), *Pseudomonas* (0.06%), *Paenibacillus* (0.05%). The proportion of *Weissella* in A4 and C4 increased greatly, more than 70% in C4, indicating that *Weissella* played a greater role in the later stage of composting. It should be noted that *Weissella* also belonged to *Firmicutes*. To this day, 19 species of *Weissella* have been identified, such as *W. beninensis, W. ceti, W. cibaria, W. confuse* etc(47,48)^]^. Among them, *W confuse* and *W cibaria* have the potential of anticancer, anti-inflammatory, antibacterial, antifungal and enhancing immunity (49,50,51), which may be beneficial to the synergistic effect of *Pediococcus* and *Weissella* in the middle and later stage of composting. The relative abundance of bacteria genera of Stage 5 was as follows, A5: *Pediococcus* (47.51%), *Weissella* (20.21%), *Bacillus* (2.60%), *Leuconostoc* (2.25%), *Bacteroides* (1.25%); C5: *Weissella* (47.99%), *Leuconostoc* (3.86%), and added bacteria of *Bacillus* (0.20%), *Pseudomonas* (0.11%), *Paenibacillus*(0.01%),and also some Chloroplast (42.61%). Both A5 and C5 were the state of natural storage for 10 days after the end of composting. In A5, *Pediococcus* still occupied the dominant position, followed by *Weissella*. In C5 period, the proportion of *Weissella* decreased significantly, and a large proportion of Chloroplast (42.61%) appeared, which may be caused by incomplete degradation of vegetable wastes. Chloroplast has appeared in the composting components since Stage 3, and always occupies a certain proportion during stage3-stage5, indicating that the microbial bacteria is still relatively limited in the degradation of vegetable waste. In the future composting process, strains with strong cellulose degrading ability can be added and their comprehensive degradation effect need to be determined. The proportion of added bacteria was maintained at a low level during C1-C5, indicating that the added bacteria were not the dominant strains in the process of composting, and it is possible that the way they affect the composting process is to change the relative abundance of dominant bacteria in different stage of composting(Fig. S1).

**Figure 4.**
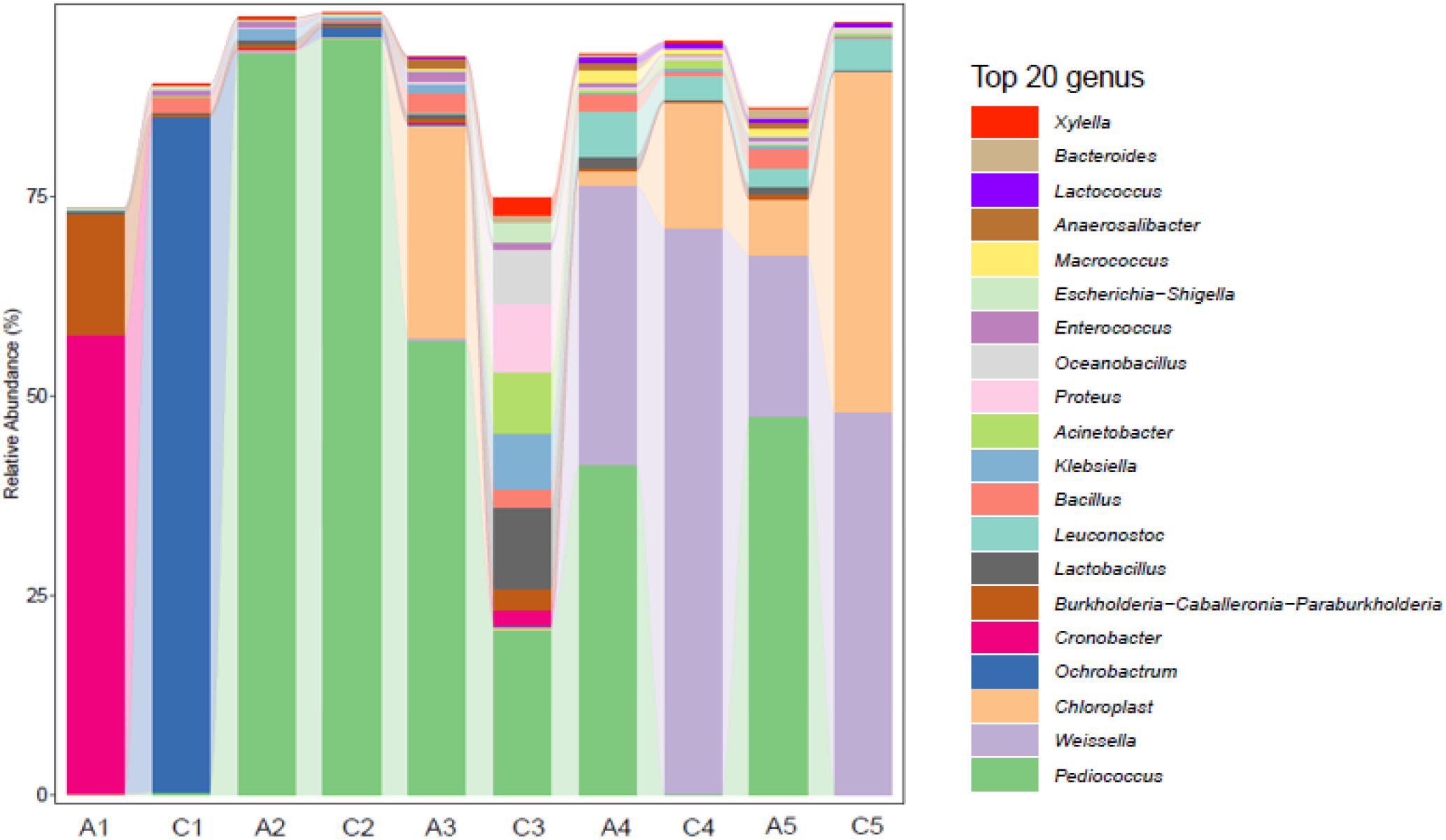
Bacteria composition at different stages of genus classification

### Analysis of Beta diversity

Through PCoA we can study the similarity and difference of data. Fig. 5 shows the PCoA results of different samples in genus level. It can be seen from Fig. 5 that PC1 is the first principal coordinate component, and its contribution to the representativeness of the total bacteria is 49.4%; PC2 is the second principal component, and its contribution to the representativeness of the total bacteria is 26.6%; for the PC1-2 analysis of the principal component, A1 and C1 are farther apart on the coordinate axis, indicating that the added bacteria may have a significant impact on the composition of the microbial community at the early stage of the HFW composting process. A2 and C2 are almost at the same position, indicating that the microbial community composition of A2 and C2 at the genus level is almost the same at this stage, and that the added bacteria species at this stage has little effect on the microbial community. A4 and C4 are far apart on the coordinate axis, indicating that the groups A and C have a large difference in the composting process in the later stage. The added bacteria may affect the proportion of dominant bacteria in different stages of the composting process to improve the composting rate and other critical factors.

**Figure 5.**
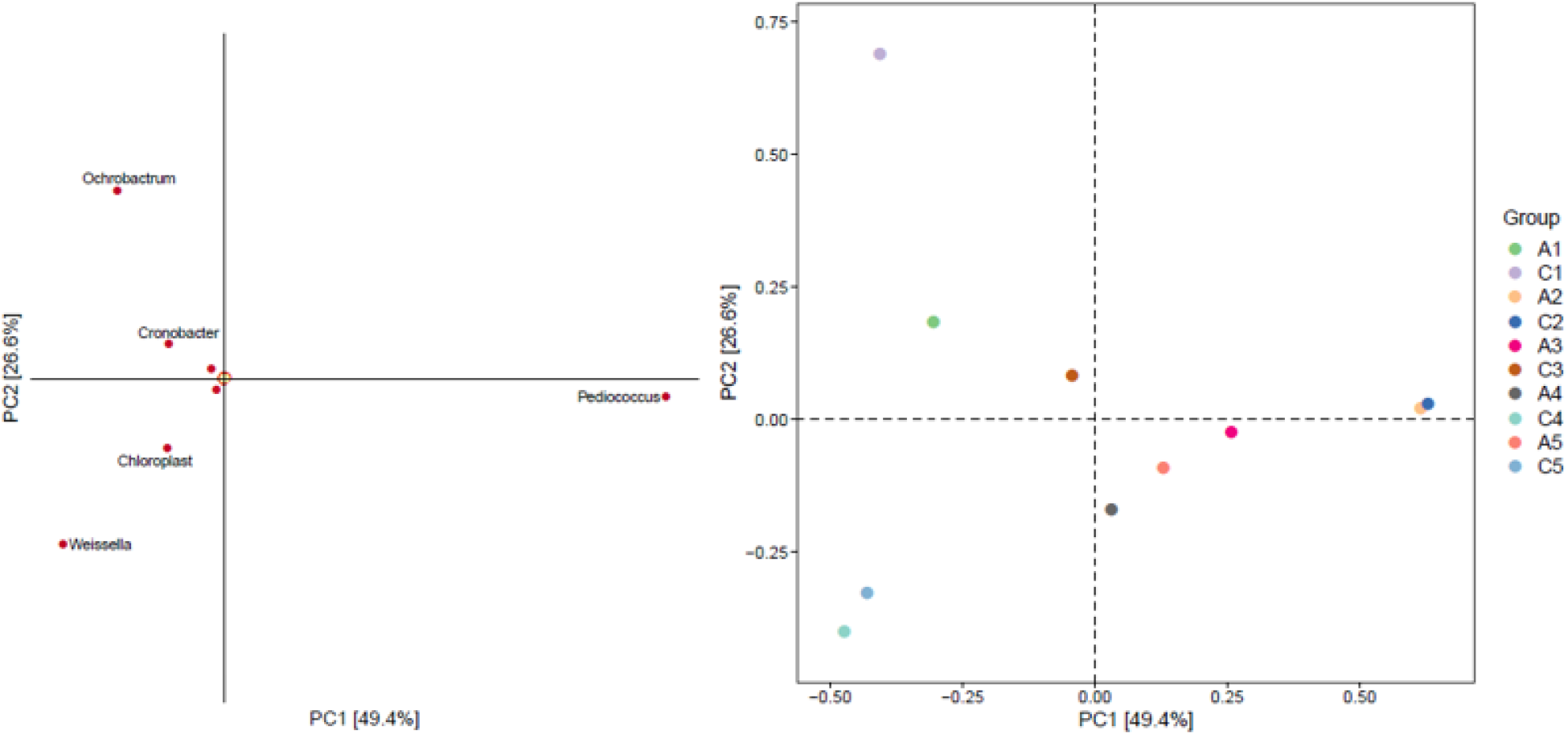
Analysis of PCoA in different samples

## CONCLUSION

The above analysis shows that at phylum level, *Firmicutes*, *Proteobacteria* and *Cyanobacteria* accounted for the largest proportion during the composting process of group A and C. At genus level, *Pediococcus* and *Weissella* were the dominant bacteria of group A and C, and both of them belonged to *Firmicutes*. The abundance of the added bacteria in the composting process of group C is always at a low level, indicating that the added bacteria may affect the composting process by changing the proportion of the dominant strains. The relative abundance of pathogenic bacteria decreased to a very low level after *Pediococcus* and *Weissella* gradually increased and occupied a dominant position in the middle stage of the two groups, indicating that *Pediococcus* and *Weissella* might play a certain inhibitory role on pathogenic bacteria. *Weissella* is the dominant bacteria in group C, and the final degradation rate of group C is higher than that of group A, indicating that *Weissella* may play a positive role in improving the degradation efficiency. In C1, the *Ochrobactrum* occupied a considerable proportion and the smell during this stage is obviously better than that of group A, indicating the *Ochrobactrum* may have an active impact on the removal of peculiar smell. Group A performed better in stabilizing the microbial community in the later stage and the *Pediococcus* took a rather high proportion during the composting process of group A, indicating that *Pediococcus* have the ability of maintaining system stable and prolonging the composting cycle.

In conclusion, the microbial communities of the two groups during the composting process of HFW have a high degree of diversity. There are not only differences in species distribution and relative abundance but also common core bacteria between them. In this paper, the study of microbial community changes in the composting process of HFW is helpful to explore the microbial resource pool, deepen the understanding of the composting process of HFW, and is of great significance for the further harmless resource treatment of HFW.

## MATERIALS AND METHODS

### Materials

The HFW was collected from the dining hall of Ningbo Tech University; The added bacteria are: B2 (*Bacillus subtilis*), F4 (*Pseudomonas aeruginosa*), 303 (*Paenibacillus lautus*), 792 (*Bacillus thuringiensis*), which are all preserved in our laboratory; Sawdust; LB broth medium; 75510-019Agarose (Invitrogen); Non-fat milk powder; NEB M0491L Q5^®^ High-Fidelity DNA Polymerase (New England Biolabs (NEB); P7598 Quant-iT PicoGreen dsDNA Assay Kit (Invitrogen); DL2000 Marker (Takara); AM9870 TAE (Invitrogen).

### Equipment

Small HFW treatment machine (Midea Group); Freeze dryer; pH meter; ABI 2720 PCR amplify instrument (ABI); FLX800T type enzyme labelling instrument (BioTek Proton Instrument Co., Ltd.); DYY-6C electrophoresis apparatus (Beijing 61 Biological Technology Co., Ltd.); Microplate reader (BioTek, FLx800); BG-gdsAUTO (130) gel imaging system (Beijing Bai Jing Biotechnology Co., Ltd.)

### Preparation of bacteria freeze-dried powder

These added bacteria which was preserved on slant culture medium were inoculated 200 ml LB medium (sterilized at 121 ° C for 20 min), and then cultured in a constant shaker at 37 ° C, 200 rpm for 24 h. After culture, the broth was centrifugated at 4 ° C for 15 min at 8000 rpm and the supernatant was discarded, and the bacteria precipitate were thoroughly mixed with 5-10 ml of non-fat milk (sterilized at 100 ° C for 10 min) and poured into the plate. After sealing the plastic wrap (poking several small holes) and pre freezing at - 4 ° C for 2 h, the plate was freeze-dried using a freeze dryer for 18 h, and then distributed into 10 ml EP tube (sterilized at 121 ° C for 20 min) and put into 4 ° C freezer for further use (about 5 g of freeze-dried powder can be made for every 200 ml of broth, the viable bacterial count of freeze-dried powder was about 10^9^ CFU/ g).

### Experimental grouping and treatment

This experiment was divided into two groups: group A (natural composting) and group C (composting with added bacteria), and the composting process was carried out in the HFW treatment machine. 500g sawdust was added at the beginning of composting of two groups. In group C, 5g freeze-dried powder with four kinds of added bacteria was added, and 500g ~ 600g HFW was added at 1 pm every day. The residue in the machine was sampled every two days to determine the number of viable bacteria and the pH value, and 20g sample was stored in the refrigerator at - 70 ° C for subsequent 16S rRNA sequencing analysis. The composting process sustained 20 days, and after that the composting residue was naturally stored for 10 days. The composting samples on the 1st, 7th, 15th, 20th and 30th days were taken for 16S rRNA sequencing analysis, named stage1-stage5. At the end of experiment, the weight of composting residue was measured and the degradation rate was calculated. The degradation rate can be calculated according to the following formula:

Degradation rate (%)= (X_1_-X_2_+X_3_)/X_1_
X_1_—Total weight of HFW added (g)
X_2_—Weight of composting residue in the machine (g)
X_3_—Weight of sawdust added (g)

### High throughput sequencing of 16S rRNA

According to the experience, the most suitable method of total DNA extraction was utilized for different sources of microbiome samples. At the same time, DNA was quantified by Nanodrop and the quality of DNA was detected by 1.2% agarose gel electrophoresis.

Generally, the target sequences, such as ribosomal RNA or specific gene fragments, which can reflect the composition and diversity of microbial community, are used to design primers according to the V3 ~ V4 region of 16SrRNA gene, and the specific barcode sequence was added to amplify the rRNA gene variable region (single or continuous multiple) or specific gene fragment by PCR.

The PCR amplification adopts PFU high fidelity DNA polymerase (TransGen Biotech), and the number of amplification cycles were strictly controlled, so as to make the number of cycles as low as possible and ensure the same amplification conditions of the same batch of samples. At the same time, a negative control was set, which can detect environmental, reagent and other microbial contamination, and any negative control amplification samples with bands can not be used for subsequent experiments. The related information are shown in table 4.

**Table 4:**
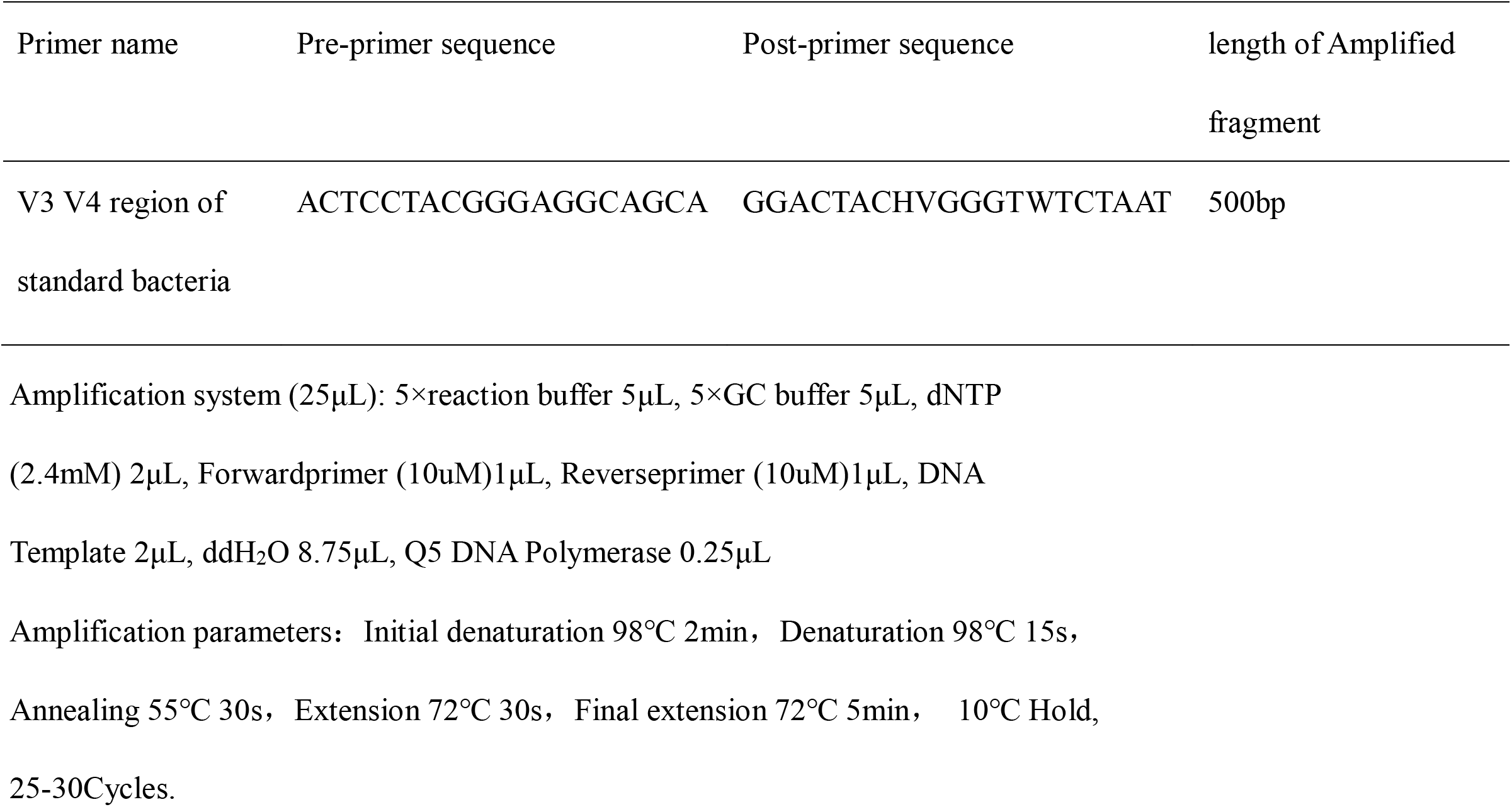
Primer information of PCR amplification

### Fluorescence quantification of amplification products

The PCR products were quantified by fluorescent reagent quant (Quant-iT PicoGreen dsDNA Assay Kit), and the quantitative instrument is Microplate reader (BioTek, FLx800). According to the results of fluorescent quantitative analysis, the samples were mixed according to the corresponding proportion according to the sequencing requirements of each sample.

### Construction and sequencing of Illumina Library

Illumina high-throughput sequencing and bioinformatics analysis were performed with the assistance of Personalbio Biotechnology Co. Ltd. (Shanghai).

## ACKNOWLEDGMENTS

We would like to thank the staff working at the dining hall of Ningbo Tech University for kindly offering us the household food waste. In addition, a particular thanks goes to Personalbio Biotechnology Co. Ltd. (Shanghai) for sample testing and chart making. This work was financially supported by Zhejiang Provincial Department of Education (item Number: Y201941839).

We declare that we have no competing interests.

